# Pollen source richness may be a poor predictor of bumblebee colony growth

**DOI:** 10.1101/2020.11.30.403444

**Authors:** Cecylia M. Watrobska, Ana Ramos Rodrigues, Andres N. Arce, Jessica Clarke, Richard J. Gill

**Affiliations:** Department of Life Sciences, Imperial College London, Silwood Park campus, Buckhurst Road, Ascot, Berkshire, SL5 7PY, UK; Current address: Department of Biological Sciences, Royal Holloway University of London, Egham, TW20 0EX, UK; Current address: Department of Biology, Lund University, SE-223 62 Lund, Sweden; Current address: Department of Life Sciences, University of Suffolk, Waterfront Building, 19 Neptune, Quay, Ipswich, IP4 1QJ, UK

**Keywords:** social bees, pollen diversity, workers, pupae, land use, monoculture, agriculture, insect pollinator, foraging

## Abstract

1. Agricultural intensification has drastically altered foraging landscapes for bees, with large-scale crop monocultures associated with floral diversity loss.
2. Research on bumblebees and honeybees has shown individuals feeding on pollen from a low richness of floral sources can experience negative impacts on health and longevity relative to higher pollen source richness of similar protein concentrations. Florally rich landscapes are thus generally assumed to better support social bees. Yet, little is known about whether the effects of reduced pollen source richness can be mitigated by feeding on pollen with higher crude protein concentration, and importantly how variation in diet affects whole colony growth, rearing decisions and sexual production.
3. Studying queen-right bumblebee (*Bombus terrestris*) colonies, we monitored colony development under polyfloral pollen diet or monofloral pollen diet with 1.5-1.8 times higher crude protein concentration.
4. Over six weeks, we found monofloral colonies performed better for all measures, with no apparent long-term effects on colony mass or worker production, and a higher number of pupae in monofloral colonies at the end of the experiment. Unexpectedly, polyfloral colonies showed higher mortality, and little evidence of any strategy to counteract the effects of reduced protein; with fewer and lower mass workers being reared, and males showing a similar trend.
5. Our findings i) provide well-needed daily growth dynamics of queenright colonies under varied diets, and ii) support the view that pollen protein content in the foraging landscape rather than floral species richness per se is likely a key driver of colony success.

## INTRODUCTION

Bees are essential insect pollinators of many wild flowers and crops, making reported declines an issue of global importance (Biesmeijer et al. 2006; Williams and Osborne 2009; Cameron et al. 2011; Burkle et al. 2013; Carvalheiro et al. 2013; Nieto et al. 2014; Gill et al. 2016). The emergence of agricultural land-use has been implicated as a contributing driver (e.g. Ollerton et al. 2014; De Palma et al. 2015; Samuelson et al. 2018; Powney et al. 2019). Large scale crop monocultures leading to fragmentation and loss of wild floral resources (Carvell et al. 2006; Goulson et al. 2015) are thought to have degraded the ‘nutritional landscape’ by lowering nectar and pollen availability to bees (Kleijn and Raemakers 2008; Scheper et al. 2014; Baude et al. 2016). Increased cultivation of pollinator-dependent crops since the 1960’s may therefore be encouraging news by helping to nutritionally subsidise wildflower losses (Westphal et al. 2003; Aizen et al. 2008). However, having large swathes of single plant species / varieties can reduce the diversity of florally sourced nectar and pollen available (Vaudo et al. 2015) leading to agri-environment schemes promoting management of diverse floral resources to support bees (Carvell et al. 2007; Pe’er et al. 2014; Gill et al. 2016). But, is focusing on floral diversity the most important criterion when informing such schemes? For instance, it may also be important to understand how the amount and/or quality of nectar and pollen provisioned in these floral habitats contributes to bee reproductive success (Dicks et al. 2015; Wood et al. 2015; Hicks et al. 2016; Moerman et al. 2017).

For bumblebees, floral pollen is the exclusive protein source needed for colony growth (Sutcliffe and Plowright 1988), influencing the number and size (mass) of reared individuals, both of which have a positive feedback on future pollen income (Duchateau and Velthuis 1988; Kitaoka and Nieh 2009). Under natural settings, however, there is a limit to the amount of pollen that can be brought back to the colony because workers are restricted by foraging ranges, weather conditions and temporal food resource gaps (Gill et al. 2016; Arce et al. 2017; Becher et al. 2018; Timberlake et al. 2019). Landscapes dominated by floral species possessing pollen of high crude protein may therefore be beneficial and help to counteract such constraints. Indeed, studies of bumblebee micro-colonies (a subset of colony workers kept together without a queen) have shown that pollen from one plant species can be associated with increased growth compared with provision from another (Ribeiro et al. 1996; Génissel et al. 2002; Tasei and Aupinel 2008b; Vanderplanck et al. 2014; Baloglu and Gurel 2015; Moerman et al. 2015, 2017; Roger et al. 2017; Leza et al. 2018). However, feeding on pollen from a monofloral source may come at a cost of losing pollen nutritional diversity (Roulston et al. 2000; Brodschneider and Crailsheim 2010; Di Pasquale et al. 2013; Vanderplanck et al. 2014), with any reduction in floral species richness potentially leading to deficiencies in essential nutrients (Vaudo et al. 2015).

An increase in the diversity of florally sourced pollen has been shown to be associated with improved individual bee condition, such as longevity (Standifer 1967; Schmidt et al. 1995; Wang et al. 2014) and immune capacity (Alaux et al. 2010, 2011; Di Pasquale et al. 2013; Brunner et al. 2014), as well as benefitting a set of micro-colony parameters (e.g. egg production and larval weight; Di Pasquale et al. 2013; Dance et al. 2017). However, when studying the effect of diet on bumblebee colony growth rates, we still have a limited understanding as to how the potential benefits of feeding on pollen from different floral species may be mediated by crude protein concentration (Moerman et al. 2015, 2017). Insights gained by such experimentation contribute to informing management practices when advising on floral composition of habitats to support bumblebees. A first step is to better understand comparative colony responses to provision of pollen of relative high crude protein content but of low floral source diversity, versus, provision of pollen of relative lower content but of higher diversity. Whilst pollen diversity could lead to increased worker longevity, we predict that feeding on pollen of lower crude protein content would lead to fewer and/or smaller individuals being reared; but empirical data on number-mass responses in bumblebees is limited (Herrmann et al. 2018). To address this requires monitoring of the day-by-day growth dynamics of bumblebee colonies under different diets (Tasei and Aupinel 2008a; Baloglu and Gurel 2015; Rotheray et al. 2017). To date, however, there is also a surprisingly limited amount of data on the relative long term daily dynamics of colony growth (Duchateau and Velthuis 1988).

Here we compared colony growth rates over six weeks in queen-right bumblebee *Bombus terrestris* colonies provisioned with a polyfloral pollen diet versus colonies provisioned with a monofloral pollen diet with a relatively higher crude protein concentration. By meticulously monitoring all newly eclosed bees and individual deaths we studied colony growth strategies in response to diet by looking if sequential generations of workers responded differently between the monofloral and polyfloral diet to investigate possible cohort lag effects. We further investigated whether the monofloral diet had long-term impacts by: i) impacting the later stages of colony development; ii) influencing colony decisions on the number-mass trade-off when rearing individuals; iii) altering mortality rate. Starting with small established queen-right colonies, this experiment measured colony food consumption, colony weight gain, worker and male production, mass of reared individuals, worker mortality, and number of pupae at the end of the six-week period.

## METHODS

### Bumblebee colonies

Twenty-four colonies were ordered from the commercial supplier Biobest NV (Belgium) and distributed by Agralan Ltd (UK). Colonies were housed inside a self-contained and ventilated plastic nest box (25×20×13 cm) for the six-week experiment (42 days). On arrival the sugar solution reservoir and pollen patty provided with the colonies were removed, and all colonies were placed in an environmentally controlled room (23°C, 60% humidity) under continual red light. Twenty-four hours prior to start of the experiment (also 24hrs after arrival) each colony was provisioned with a feeder containing 25mL of 40% sucrose solution, checked for an active queen, and the number of pupae and workers counted.

Two experimental replicates (ERs) were conducted: i) for ER1, colonies arrived at the approximate size requested with monofloral assigned colonies (n=6 colonies) having a mean (±s.e.m.) of 34.2±3.9 pupae and 24.7±2.0 workers, and polyfloral colonies (n=6) having 26.8±2.8 pupae and 22.8±1.2 workers; ii) for ER2, colonies arrived slightly larger than requested with monofloral colonies (n=6) having 42.2±5.7 pupae and 47.0±4.8 workers, and polyfloral colonies (n=6) having 37.2±7.9 pupae and 50.5 ± 1.6 workers. Therefore, workers from 11 of the ER2 colonies were culled (random removal) to reduce worker number per colony to 35 (size of the smallest colony; Table S1). For each ER, colonies were assigned to treatments by ranking first the number of workers present in the colony on arrival (prior to culling for ER2), which was then coupled with a count estimate of colony pupal number. The sum of these two ranks was then determined with each colony paired with its closest consecutive rank and assigned randomly to either the monofloral or polyfloral treatment. We found no significant difference in worker numbers between monofloral and polyfloral assigned colonies (GLM: workers: z=0.34, p=0.74) or initial colony mass (LM: t=1.38, p=0.18). Based on our estimated pupal counts there was however a significantly lower number of pupae in polyfloral colonies (meanΔ=-16%; GLM: z=-2.55, p=0.011; Table S2), which was considered when running our statistical analyses.

### Feeding regime and pollen diets

During the experiment, colonies were provided 40% sucrose solution in a gravity feeder alongside the respective pollen diet inside a 55mm diameter (12mm deep) Petri dish, three times per week (Monday, Wednesday & Friday; colonies fed from days 1-38). For sucrose and pollen provision, we incremented the standardised volume/mass as the experiment progressed (see Table S3 for set amounts). Our choice to provision limited amounts rather than *ad-libitum*, was to simulate a more realistic scenario of colonies being constrained by physical access and availability in the field (see Introduction). Honeybee collected pollen (supplied by Agralan) was used for making the pollen diets and was stored at −20°C. A subsample was taken each time for provisioning, which ensured it had only just thawed before being provided to colonies. Each time the sucrose feeder and pollen diet provisions were replenished, any remaining sucrose solution and pollen were measured to the nearest 0.1mL and 1μg respectively.

Different colours of the supplied honeybee collected pollen pellets indicated pollen was from multiple flower species. Based on an established method (Alaux et al. 2010; Di Pasquale et al. 2013; Moerman et al. 2017), we separated pellets that exhibited either a distinct light purple (ER1) or navy (ER2) colour. These two colours were chosen because they were highly distinguishable from the other pollen colours, giving us confidence to reliably and consistently pick pollen of the same colour. Furthermore, preliminary analysis of pollen content showed a relatively high level of protein content compared to the other pollen types in the mix, with protein content being similar between purple and navy. The polyfloral diet for both ER1 and ER2 consisted of the remaining pollen mixed with the purple or navy pollen constituting 5% (w/w), respectively. Basing the monofloral diet on colour inspection assumed that each similarly coloured pollen ball was comprised of mostly one species of flowering plant, considering that honeybees are often florally constant during each foraging bout (Free 1963). Palynological analysis of each pollen diet to identify richness and composition of the pollen morphotypes, alongside a Bradford assay to measure crude protein content, provided support to justify our assumptions made above (see Results).

### Pollen analysis

Using a compound microscope (Labophot-2, Nikon) we determined the morphology of pollen grains from subsamples of pollen from each diet. The following protocol was undertaken to prepare samples for observation under the microscope: 1) 2g subsample of each pollen diet (ER1 monofloral, ER1 polyfloral, ER2 monofloral, ER2 polyfloral) was taken and placed in a separate 50mL falcon tube along with 20mL of deionised water. Each sample was vortexed for 1min to produce a well-mixed homogenous ‘pollen stock solution’ (at a 0.1mg/μl concentration). 2) A ‘fuchsine and spore solution’ was previously prepared: with a 25mg/ml ‘spore suspension stock’ first being prepared by dissolving *Lycopodium* spores in deionised water; followed by adding a volume of ‘fuchsine and Ethanol stain solution’ (0.01g/ml) to obtain a final fuchsine and spore solution at a concentration of 1.25mg/ml. Note: the highly consistent diameter size of *Lycopodium* spores (33μm) were integrated into the sample to provide a standardised size marker, and the fuchsine dye was used to increase contrast and so enhance identification of different pollen grain features (Sawyer and Pickard 2006). 3) From each pollen stock solution 10μl was aliquoted into separate wells of a multi-well microplate with an additional 90μl of the ‘fuchsine and spore solution’ added to each. 4) This 100μl sample was left to stain for 5mins. 5) For each diet, 3μl of the 100μl sample was pipetted out on to a 75×25mm glass microscope slide to provide a sample spot which was repeated four times. 6) Once the sample spots were dry, 10μl of warmed 80% glycerol was pipetted on top of the sample and an 18×18mm cover slip was placed over the top.

Classification of the pollen grain morphotypes was conducted from photographic images produced by fitting a GX-CAM digital camera (GXCam-5, IS500, 5MP; GT Vision Ltd) to the microscope. Images were captured at x400 magnification at a resolution of 2592×1944 pixels, and manually adjusted exposure using the GT Vision software. For each of the four sample spots per pollen diet, we obtained images of five randomly selected microscopic fields-of-view (total=20 images; each field of view=218.7mm^2^). Morphotypes were described using a combination of seven main characters: i) size; ii) shape; iii) thickness and structure of the exine (outer layer); iv) ornamentation of the exine; v) observable apertures and the number; vi) aperture type; and vii) level of staining (Sawyer and Pickard 2006; Table S4). As morphotype identification progressed, each morphotype was attributed a unique number and added to a reference picture library compiled to aid in classifying all future pollen morphotypes (Figure S1). For each morphotype, we counted the number of pollen grains observed across the 20 images. Taking three representative pollen grains per morphotype, we measured the width at the widest point of the grain and took the mean value. To gain a relative proportion of each morphotype per diet, we took morphotype abundance and divided by the mean grain width, to control for many smaller grains taking up the same space as fewer large grains.

We investigated what the relative differences were in crude protein content between the pollen diets by conducting a Bradford protein assay (Bradford 1976). We first prepared the samples for analysis by washing 1g of each pollen diet in 10ml of ultrapure Milli-Q water and vortexed the samples until the pellets had dissipated and the pollen grains had entered suspension. The suspension was centrifuged at 6000rpm for 5mins and the supernatant was discarded. The remaining pollen was left to dry overnight at room temperature under a fume hood. We then placed between 97-100mg of pollen into individual 2ml homogenisation tubes, containing 600mg of 1mm ceramic homogenisation beads. The tubes were placed at −80°C for 40mins prior to homogenisation at 30MHz for 5mins (Qiagen Tissue Lyser II). After the initial homogenisation, we added 0.1M NaOH to create approximately 1ml of a 0.1mg/ml pollen lysate which was heated to 100°C for 5mins before re-homogenising the lysate for a further 3mins. We then conducted the Bradford assay using the Bio-Rad Protein Assay Kit microassay 300μL microplate protocol using bovine serum albumen as the protein standard (Bio-Rad Laboratories). Because of the high protein concentration of the pollen, we conducted the assay on the lysate after serial dilution in Mili-Q water by a factor of 10^-3^. We assayed each pollen sample in triplicate wells containing 40μl Bradford reagent mixed with 160μl of pre-prepared BSA protein standards (0-80μl/ml). The plates were incubated at room temperature for 5mins, followed by 30secs of shaking, before reading the absorbance at 595nm (spectrophotometer, Synergy HT, Bioteck). Protein concentrations were calculated using linear regression analysis from the protein standards.

### Colony monitoring

Prior to the start of the experiment all workers per colony were tagged with a unique colour and number Opalith tag. Between experimental days 1–35, colonies were checked daily (Monday-Friday) and any newly eclosed workers were tagged allowing us to estimate the age of each bee. Bees were not tagged between days 36-42, and so any untagged bees found at the end of the experiment were determined to have eclosed during this last week. Tagging involved removing a newly eclosed bee from a colony using forceps, placing inside a marking cage and applying the tag using superglue. The bee was then allowed to rest for 5mins in an individual holding container, after which the bee was placed back into the colony. Any dead bees found inside the colony during the tagging process were removed, tag (if present) noted and placed inside an individual 2 ml tube and frozen at −20°C.

On day-1 each colony was weighed (AE Adam^®^, model PGW1502e, accuracy±0.01g) and a repeated measure of colony mass was taken every seven days with the mass of the plastic box subtracted. Colony mass therefore constituted all individuals, wax nest, brood and any pollen and/or nectar stores. On day-42 all colonies were sacrificed by placing the colony box in a freezer (−20°C), and after 24hrs each colony was dissected and number of workers, gynes (newly produced queens), males (drones) and pupae were counted, and mass of the nest structure weighed. The wet mass per worker was taken, and additionally the mean wet mass per pupae was calculated by taking the total mass and dividing by the total number of individuals. Whilst we considered the number of individuals that had eclosed and their mass for each week in our analyses, for dead individuals we only considered the total dead by the end of the experiment for two reasons: i) if an individual dies underneath the brood or in an inconspicuous place then the date of death is difficult to accurately determine; ii) dead individuals decompose resulting in inaccurate measures of mass.

### Statistical analysis

For all analyses *diet* was considered as the predictor. Error structures were adjusted according to the response variables, with linear mixed effects models (LMER) using a gaussian error distribution used for continuous response variables (consumption, individual mass & colony mass) and generalised linear mixed effects models (GLMER) with a Poisson distribution (link function = “log”) for count variables (numbers of adult individuals & pupae). To analyse total worker mortality by the end of the experiment, we used a GLMER with binomial error distribution. Experimental replicates (*block*) were incorporated as a fixed factor in the models. For analysis of repeated weekly measures (colony mass, weekly consumption, weekly worker eclosion, and weekly worker mass) we: i) included observation *week* nested in *colony* as a random factor to account for temporal pseudoreplication of colony measures; and ii) ran each as a linear and second order polynomial (quadratic) model, choosing the best fitting model based on AIC values. When considering mass of individuals, male wet mass was log10 transformed in response to non-normal distribution of the data. For models of worker production over time and end-point total number between diets, we compared models with and without colony starting pupae number as an explanatory covariate. Statistical outputs stated in the main text are from models that did not include starting pupal number, as outputs showed the same significant patterns (Tables S9, S10). All analyses were conducted in R v3.5.1 (R Core Team 2017) with packages ‘lme4’ v1.1.21 for mixed effects models (Bates et al. 2015), ‘lmerTest’ v3.1.0 to calculate corresponding p-values (Kuznetsova et al. 2017) and ‘ggplot2’ for data visualisation (Wickham 2016).

## RESULTS

### Pollen analysis

We identified a total of 16 pollen morphotypes in the supplied honeybee collected pollen (Figure S1), with ER1 & ER2 polyfloral diets consisting of 14 and 13 morphotypes, respectively. Nine of the 14 morphotypes (ER1) constituted 2.2–37.8% composition of the pollen diet based on relative pollen grain counts (Shannon-Weiner Index (H) of pollen community=1.90). Eleven of the 13 morphotypes (ER2) constituted 2.2-19.6% (H=2.23). ER1 monofloral diet consisted of five of the total 16 morphotypes with one of these morphotypes (morph A) constituting 96.8% of the diet (H=0.17). For the ER2 monofloral diet, a single morphotype (morph F) constituted 100% of the diet (Table S5). The Bradford assay, based on the mean crude protein content of the pollen lysate (mg/ml), showed the monofloral diets had a 1.8 and 1.5-fold higher relative protein concentration than the polyfloral for diets compared in ER1 and ER2 respectively (mean value: ER1=59.4 vs 32.9mg/ml; ER2=63.2 vs 41.5mg/ml; Table S6).

### Food consumption

Consumption of pollen and sucrose solution showed a parabolic type trend, increasing over the first 4 to 5 weeks of the experiment and decreasing in weeks 5 and 6 (LMER: *week*^2^: pollen, t=-5.56, p<0.001; sucrose, t=-4.54, p<0.001). For pollen consumption, there was no significant difference between monofloral and polyfloral colonies (*treatment*^2^: t=0.29, p=0.77) and this was consistent over the duration of the experiment (*treatment*week*^2^: t=0.04, p=0.97; Figure S2, Table S7), with colonies consuming a median (IQR) of 95.1% (77.0-97.5) versus 88.5% (76.6-99.0) of the total pollen provided, respectively. For sucrose consumption, monofloral consumed significantly more than polyfloral colonies (*treatment^2^:* t=3.92, p=0.001), which remained relatively consistent over the course of the experiment (*treatment*week*^2^: t=0.96, p=0.34; Figure S3, Table S7), with 88.1% (82.4 - 94.5) versus 76.6% (69.7 – 82.3) of the total provisioned volume of sucrose consumed.

We then analysed relative consumption by taking the amount consumed per gram of colony mass (total mass (pollen) or volume (sucrose) divided by colony mass each week). Relative pollen and sucrose consumption significantly decreased over the consecutive weeks (LMER: *week:* t=-2.79, p=0.011 & t=-8.21, p<0.001, respectively), but this response did not significantly differ between monofloral and polyfloral colonies (*treatment:* t=0.18, p=0.86 and t=-0.75, p=0.46) and did not change over time (*treatment*week:* t=-0.13, p=0.90 and t=-0.51, p=0.62; Figure S2-3, Table S7).

### Colony Growth

Over the course of the experiment our model suggests that the weekly colony growth rate (positive linear term) was significantly higher in monofloral relative to polyfloral colonies (LMER: *treatment*week:* t=-3.23, p=0.004, Figure 1A, Table S8). Accordingly, mean polyfloral colony mass was 18.84g (−22.1%) and 11.34g (−9.8%) lower relative to monofloral colonies in ER1 and ER2 respectively (Figure 1B). Cumulative numbers of weekly worker eclosure did not differ significantly between monofloral and polyfloral colonies (GLMER: *treatment:* t=1.64, p=0.10; Table S9). However, there were consistent negative model estimates for polyfloral colonies, and when comparing endpoint data (cumulative total of workers produced over the whole experiment), we found monofloral colonies had reared a significantly higher number of workers compared with polyfloral colonies (GLMER: z=-6.17, p<0.001; Figure 2A); an effect found when also considering colony starting pupae number (z=-4.22, p<0.001; Table S10).

**Figure 1:**
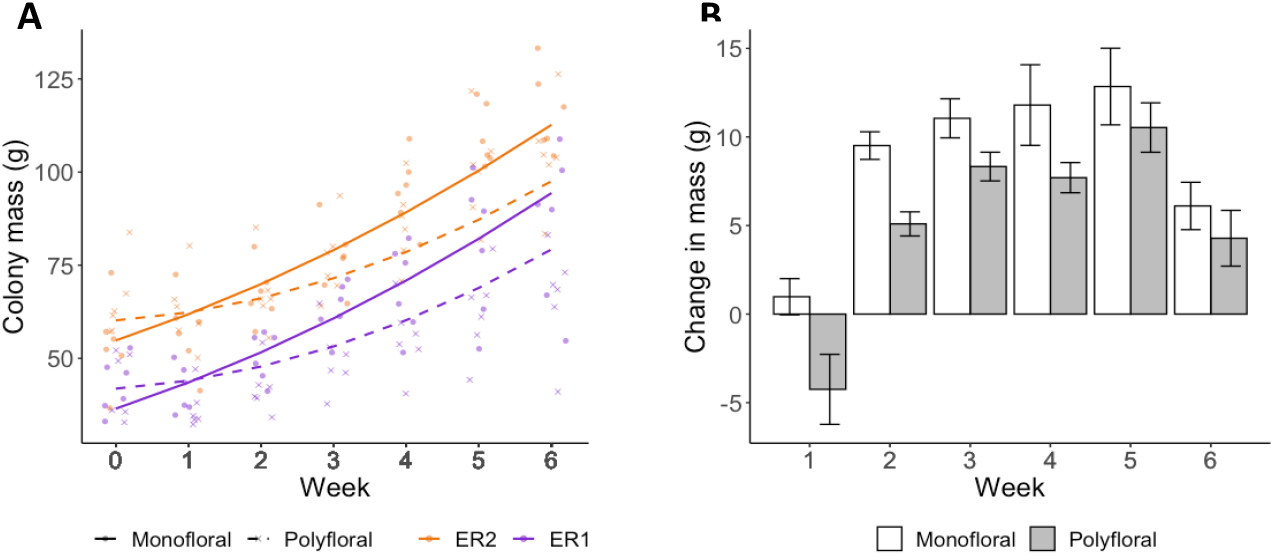
Colony growth rates. Weekly measures of colony mass (grams) for the colonies provisioned a monofloral (n=12) and polyfloral (n=12) diet. **A)** Scatter plot of cumulative colony growth rate with fitted lines representing the LMER estimates (monofloral = solid line/dot symbol; polyfloral = dashed line/cross symbol; ER1 = purple lines/symbols; ER2 = orange lines/symbols); **B)** Histogram showing mean (±s.e.m.) colony weight change (grams) by the end of each week based on raw colony mass measures.

**Figure 2:**
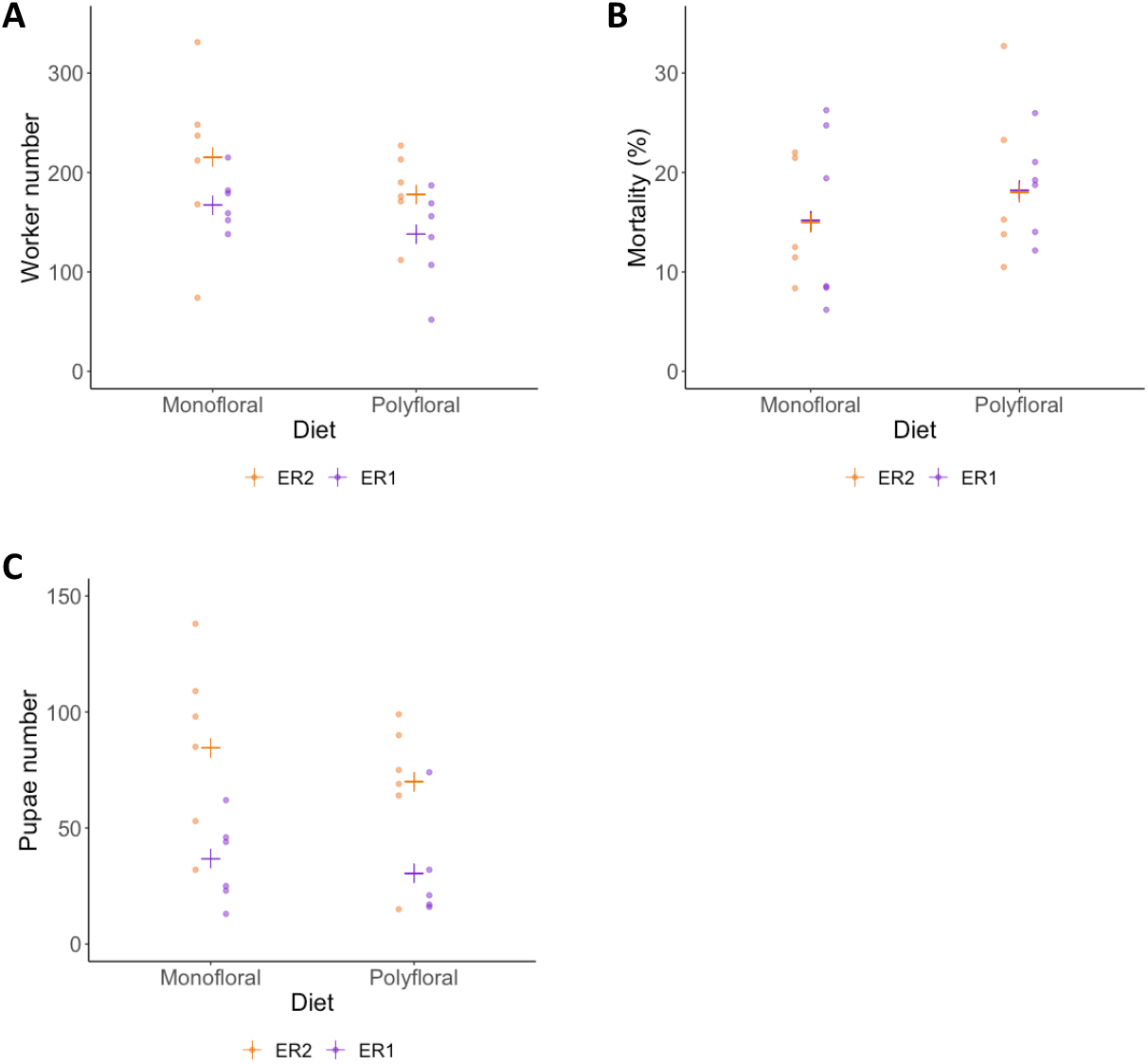
End of experiment comparisons of A) total adult workers reared (worker production), B) percentage mortality of total adult workers, and C) total number of pupae present at end of experiment. Raw data points are shown (ER1 = purple symbols, ER2 = orange symbols), with the estimated mean (cross symbols) from back transformation of the GLMER). (N.B. for panel C the colony that had produced gyne pupae was not included).

Worker mass was taken from all workers alive at the end of the experiment for which their tag could be successfully identified (n=3,356). Overall, mean worker mass showed a decrease over the first three weeks (likely due to pollen provision being limited) but appeared to plateau over the latter three weeks of the experiment (LMER: *week*^2^: t=16.35; p<0.001; Figure 3A). Our model suggests that the rate of decreasing mass (linear term) was significantly greater in polyfloral relative to monofloral colonies (*treatment*week:* t=-2.70, p=0.013; Table S11). Indeed, when pooling all workers that eclosed in weeks 1-3, worker mass between monofloral and polyfloral colonies was similar with a mean difference of 0.06% (mean±s.e.m.: 150.1±1.6 vs 149.9±1.8mg), but workers that eclosed during weeks 4-6 in polyfloral colonies had a 17.9% lower mass compared with workers eclosed in monofloral colonies during that time period (124.5±1.2 vs 102.3±1.2mg).

**Figure 3:**
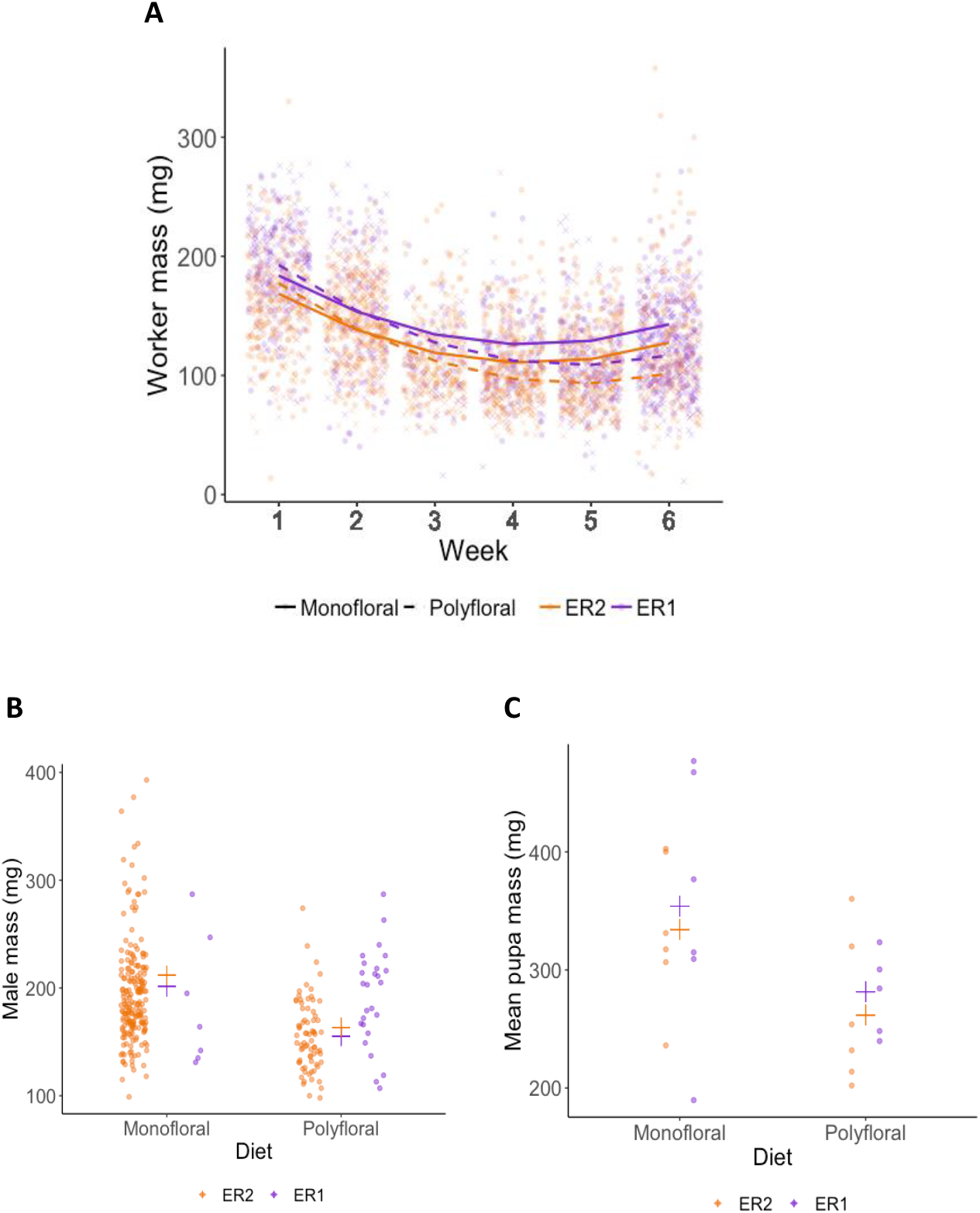
Wet mass (milligrams) of A) workers, B) males (drones), and C) pupae. **A)** Scatter plot showing mass of workers that eclosed in each week of the experiment. Fitted lines represent the LMER estimates (monofloral = solid line; polyfloral = dashed line), and the data points per week are jittered (monofloral = dot symbol; polyfloral = cross symbol; ER1 = purple lines/symbols; ER2 = orange lines/symbols). **B)** Mass of all adult males that had been produced by the end of the experiment between monofloral (n=10) and polyfloral (n=7) colonies. Jittered raw data points are shown (ER1 = purple symbols; ER2 = orange symbols), with the estimated mean from back transformation of the LMER overlaid (cross symbols). **C)** Mean pupae mass per colony present at the end of the experiment in monofloral and polyfloral colonies. Raw data points are shown (ER1 = purple symbols; ER2 = orange symbols), with the estimated mean from back transformation of the LMER overlaid (cross symbols). (N.B. for panel C the colony that had produced gyne pupae was not included).

Analysis of total worker mortality during the six-week experiment revealed lower mortality in monofloral relative to polyfloral colonies (GLMER: treatment: z=-2.79, p<0.01; Figure 2B, Table S12). Note that the mortality analysis shown here considered 22 of the 24 colonies, as one monofloral and one polyfloral colony had mortality rates of 55% and 52%, respectively, and were not considered (due to meeting model assumptions). However, analysis considering all 24 colonies showed the same significant trend; Table S12).

Males eclosed in 10 monofloral and seven polyfloral colonies (Fishers Exact Test: p=0.37). The median (range) number of males produced by colonies was higher in monofloral compared with polyfloral colonies (4 (1-108) vs 1 (1-47); Table S13), but the high ranges showed a heavy right skew in the data with a few colonies producing a particularly high number of males, making interpretation of this effect difficult. Interestingly, in parallel to our finding of lower worker mass in polyfloral colonies, we found a trend towards males of lower mass being reared by polyfloral relative to monofloral colonies (n=308, LMER: *treatment:* t=-2.10, p=0.063; Figure 3B, Table S14). No gynes were recorded to have eclosed in any colony.

Dissection of colonies at the end of the experiment revealed monofloral colonies possessed a significantly higher number of pupae relative to polyfloral colonies (GLMER: treatment: z=3.39, p<0.001; Figure 2C, Table S10 & S14). These pupae also had a significantly higher mean pupal wet mass (LMER: *treatment:* t=-2.41, p=0.026; Figure 3C, Table S15). Note that one polyfloral colony did possess one new gyne pupae (approx. 600mg), which was excluded from the above analysis.

## DISCUSSION

Our experiment simulated a hypothetical scenario whereby bee colonies had access to a set quantity of pollen resource with half the colonies having access to pollen from multiple floral species (polyfloral) and the remaining half restricted to pollen from a single species but with a 1.5-1.8 times higher relative crude protein concentration (monofloral). Over the six weeks we found that for nearly every colony growth measure, monofloral performed significantly better than polyfloral colonies. This included higher worker mass and lower mortality, which suggests that in the context of colony growth dynamics in a controlled low-challenging environment, crude protein concentration of pollen can potentially outweigh access to a higher diversity of florally sourced pollen.

### Colony development, mortality & male production

We found a higher cumulative growth rate in respect to colony mass in monofloral colonies, which was also reflected by a higher number of workers being reared by the end of the experiment. Generally, we found no clear evidence that low diversity of florally sourced pollen (monofloral) had a long-term negative effect on our colony measures (although colony mass between monofloral and polyfloral colonies appeared to show a more paralleled rate of growth in the latter couple of weeks). If diversity was beneficial in the longer-term, we might have expected to see pupae number and mean pupae mass to have been similar, if not higher, in polyfloral compared to monofloral colonies by the end of the experiment. Instead, we found polyfloral colonies to have significantly lower pupal number and mass at the end of the experiment. Considering previous findings by Dance et al. (2017) showing increased brood numbers in queenless *B. terrestris* microcolonies fed a polyfloral and not monofloral diet (although of similar crude protein content), our findings indicate that whilst bumblebee colony growth may be able to cope with reduced pollen diversity it is dependent on the ‘right’ floral species being available (Génissel et al. 2002; Tasei and Aupinel 2008b; Baloglu and Gurel 2015; Moerman et al. 2017; Leza et al. 2018).

Despite our controlled lab study imposing no intentional stress on workers, we found worker mortality to be higher in polyfloral colonies. This finding was surprising given pollen diversity has previously been shown to improve worker condition and increase longevity in honeybees (Standifer 1967; Schmidt et al. 1995; Wang et al. 2014), and considering we were able to distinguish 16 pollen morphotypes in our polyfloral diets we presume nutrient diversity will have been high (e.g. amino acids, lipids and sterols; Roulston and Cane 2000; Brodschneider and Crailsheim 2010; Di Pasquale et al. 2013). A possibility is that our monofloral sources of pollen could each represent a kind of ‘superfood’ containing all or many of the essential nutrients required for bumblebee health. Interestingly, previous studies have found large variation for individual and micro-colony level response to pollen sourced from different plant species (Ribeiro et al. 1996; Génissel et al. 2002; Tasei and Aupinel 2008b; Vanderplanck et al. 2014; Baloglu and Gurel 2015; Moerman et al. 2015, 2017; Leza et al. 2018). We should also consider that whilst mortality was lower in monofloral colonies in our controlled laboratory setup, differences may not manifest or diet effects may switch when under more challenging conditions, with workers from polyfloral colonies possibly being more resilient to stress, such as exposure to pathogens or pesticides (Di Pasquale et al. 2013; Dance et al. 2017).

The production of sexuals underpins colony fitness and changes in diet are likely to be a key determining factor, and our counts of male production per colony could be considered as a fitness proxy. A previous study by Dance *et al.* (2017) reported a monofloral diet to have a negative effect on males (although crude protein was similar with comparative polyfloral diet: 10.2 vs 12.6%) with colonies rearing fewer and lighter individuals. In contrast to this, we found no difference in the number of males reared between monofloral and polyfloral colonies, and in fact male body mass was actually higher for those reared in monofloral colonies. Considering smaller males can have a lower probability of mating (Beekman and van Stratum 1998; Amin et al. 2012; Gosterit and Gurel 2016), our findings reiterate the point that access to the right floral species, rather than floral diversity, may be vital for determining colony fitness (Regali and Rasmont 1995).

### Individual number:size trade-offs

Regarding worker production, Herrmann *et al.* (2018) showed that mean mass of workers is important for the later production of gynes (new queens) in bumblebee colonies (*Bombus impatiens*). We may therefore expect that under low food resources, such as reduced amount of absolute crude protein income, a colony may decide to alter the number of individuals it rears in order to maintain a certain average mass of reared individuals. However, we found that polyfloral colonies, that had access to a lower amount of crude protein, reared both a lower number and mass of workers - particularly in the latter two weeks of our experiment (a similar pattern was observed for reared males). Producing a smaller and lighter workforce in polyfloral colonies may have implications for colony task performance, given that larger workers appear to show better learning and foraging performances (Goulson et al. 2002; Spaethe and Weidenmüller 2002; Jandt et al. 2009; Riveros and Gronenberg 2009; Willmer and Finlayson 2014; Smith et al. 2020). The lack of any apparent compensatory strategy observed in our study is unlikely to be down to a reduced appetite, as the per capita consumption rate of both sucrose solution and pollen was not significantly different and actually higher on average in polyfloral colonies (Figures S2-3). Perhaps, therefore, the decrease in worker number and/or worker mass was not of concern to the colony with mean worker mass large enough to still contribute to the primary tasks required for colony functioning. Or, since workers could not leave the confines of our lab reared colonies, maybe there was a lack of stimulus to implement such a trade-off, given total mass of pollen income to the colony was consistent. On the other hand, the protein level in polyfloral colonies could have been low enough that any trade-off with mass would have lowered worker numbers to a point of having a substantial detrimental effect to colony function. To more formally test this, a gradient of crude protein concentrations would be needed.

### Future work and implications of our findings

Investigating the dynamics of full-sized queenright colonies is practically and financially challenging, but future work will benefit from increasing treatments and implementing a set of crossed-designed experiments to: i) undertake a combination of high and low protein concentrations with high and low pollen diversity to better identify the threshold trade-offs between pollen diversity and concentration; and ii) investigate how variation among the other pollen nutrients may compensate for a reduced diversity of florally sourced pollen (i.e. key amino acids, nitrogen content, lipids (fatty acids) and sterols (Roulston and Cane 2000; Brodschneider and Crailsheim 2010; Di Pasquale et al. 2013; Moerman et al. 2017). When considering point ii, it was interesting to find that the two different monofloral pollen species showed a consistent positive effect (with similar effect size) on colony development compared to polyfloral colonies, despite the likelihood of some aspect of nutrient composition differing between them. This reinforces the importance of high crude protein concentration for colony development. However, further studies are needed to assess the response of individuals to manipulated stress when reared under diets similar to ours. In doing so, it will improve our understanding of the relative importance of pollen diversity versus protein concentration under varied challenging scenarios (Di Pasquale et al. 2013; Leza et al. 2018).

The pollen isolated to make up the monofloral pollen diets constituted a relatively high proportion of the pollen that was ordered from the commercial supplier, and is a composition representative of that collected by honeybee foragers from commercial hives. Considering the benefit(s) reported in our study, it is highly plausible that honeybee foragers had a preference to visit this floral species or genus in their surrounding landscape, whether because it was in high abundance and/or they were attracted to it over others in the area (Aronne et al. 2012). We note that the colour and grain morphology of the purple pollen (ER1 monofloral) looks similar to pollen sourced from plants in the *Phacelia* genus, which are commonly grown in agricultural areas across Europe as either a cover crop and/or a constituent of flowering field margins (Williams and Christian 1991; Carreck and Williams 2002; Deveci and Kuvanci 2012). If our deduction is correct, we can tentatively suggest that high coverage of *Phacelia* plantings over a long enough time span and appropriate phenology could be a good food source to support colony growth of bumblebees. Indeed, bumblebee species have been shown to be able to detect and preferentially forage on floral species that provide high protein content and essential nutrients they require if available (Hanley et al. 2008; Mapalad et al. 2008; Leonhardt and Blüthgen 2012; Somme et al. 2015; Kriesell et al. 2017).

In conclusion, when designing insect pollinator and particularly bumblebee suitable landscapes, our study supports the view that we cannot necessarily assume higher floral diversity is always better in terms of supporting specific pollen dependent insects, unless we consider the nutritional content of each of the composite floral species (Wood et al. 2015; Moerman et al. 2017).

## CONTRIBUTIONS

CW, ARR, JC & RJG conducted the experiments; CW, ARR, ANA & RJG carried out the pollen and data analyses; CW & RJG wrote the paper.

## ACKNOWLEDGEMENTS

We are grateful to Dylan Smith for helping with bee husbandry, Paul Beasley for technical support and Steve Gill for advising on pollen analysis. This study was supported by a Royal Society research grant (RG130455) awarded to RJG. AA and ARR were supported by a NERC new investigator grant (NE/L00755X/1) awarded to RJG. This work contributes to the Imperial College’s Grand Challenges in Ecosystems and the Environment that supports RJG’s research.

## Supplementary Figures

**Figure S1.**
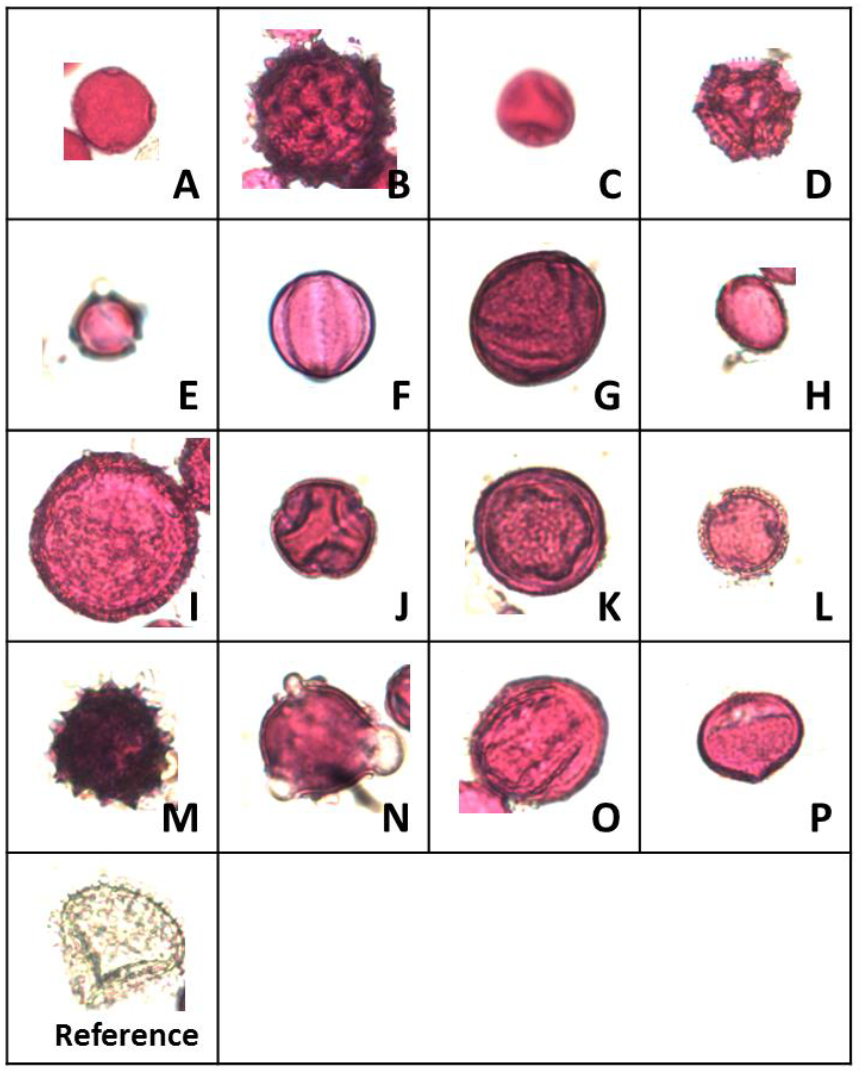
The 16 different pollen grain morphotypes identified in the honeybee collected pollen used across all four pollen diets in this experiment. Pollen was stained using a fuchsine dye and observed at ×400 magnification, with each image cropped (and hence not to scale) to show the pollen grain morphotype of interest. The reference panel shows a *Lycopodium* spores which was used as a size reference as it consistently measures 33μm at the widest point. Please see Table S5 for pollen grain diameters. Morphotype A was the dominating species in the ER1 monofloral pollen diet, and morphotype F was the dominant in ER2 monofloral diet.

**Figure S2.**
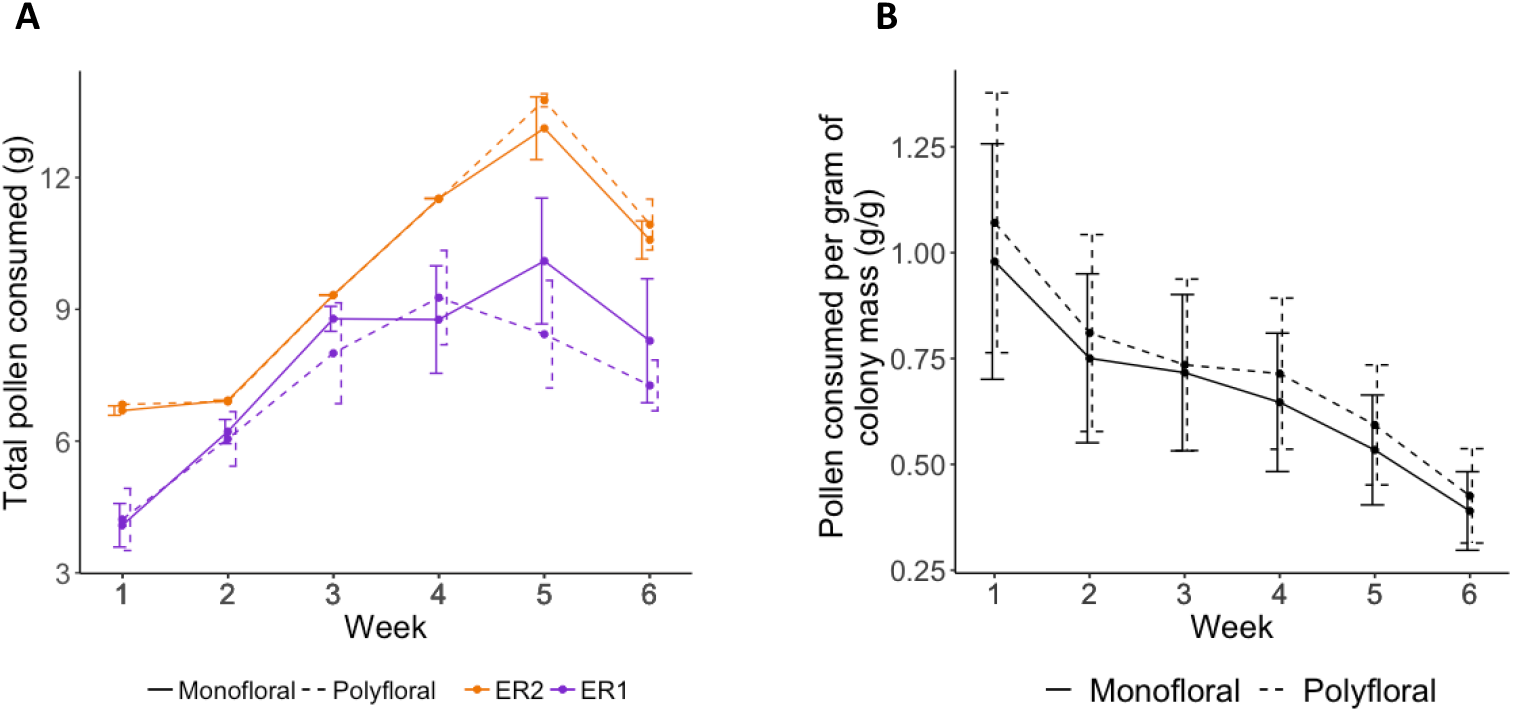
Weekly amounts of pollen consumed over the course of the experiment. Mean (± s.e.m) **A)** of the total mass (grams) of provisioned pollen consumed per colony per week (monofloral = solid line; polyfloral = dashed line; ER1 = purple; ER2 = orange); **B)** relative amounts of pollen consumed per unit of colony mass (gram per gram) for monofloral and polyforal colonies. Solid lines represent monofloral colonies and dashed line represents polyfloral colonies.

**Figure S3.**
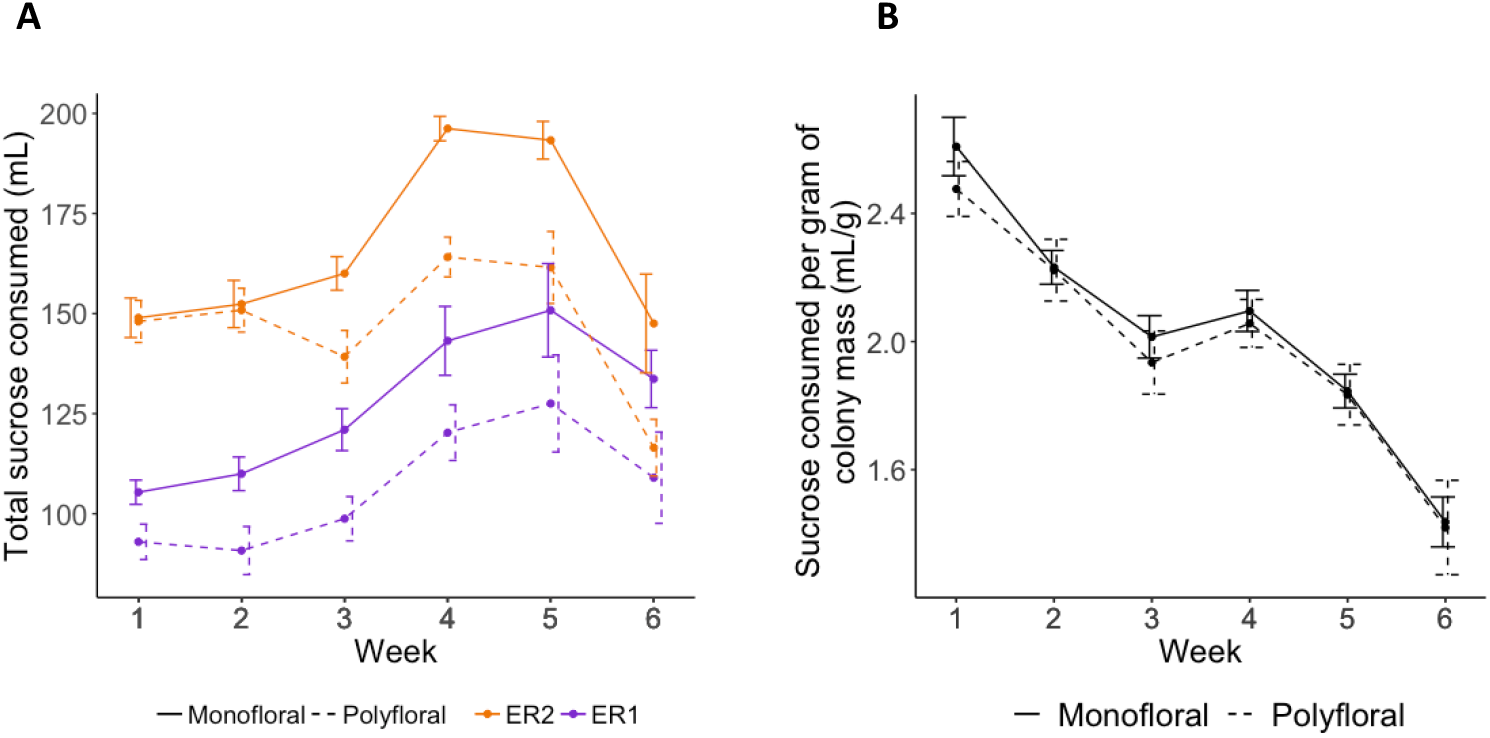
Weekly amounts of 40% sucrose solution consumed over the course of the experiment. Mean (± s.e.m) **A)** of the total volume (ml) consumed per colony per week (monofloral = solid line; polyfloral = dashed line; ER1 = purple; ER2 = orange); **B)** relative volume consumed per unit of colony mass (ml per gram) for monofloral and polyforal colonies. Solid lines represent monofloral colonies and dashed line represents polyfloral colonies.

## Supplementary Tables

**Table S1.**
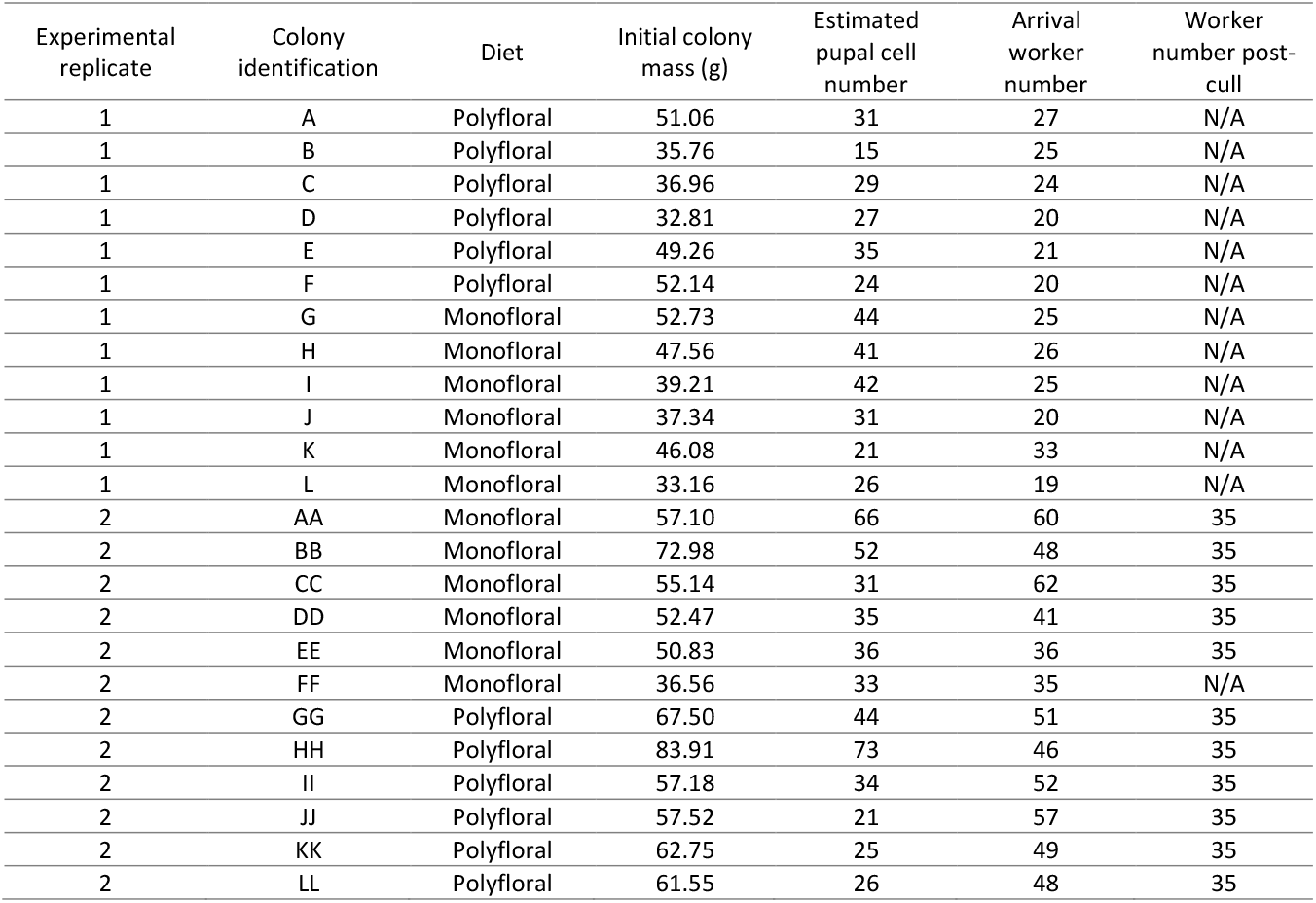
Census of each colony on arrival. ER2 colonies arrived larger than expected, and so eleven of the twelve colonies were culled (by random removal of workers) to the size of the twelfth smallest colony.

**Table S2.** Statistical outputs from linear and generalised linear models (GLM & LM) in R when analysing the difference in the a) number of workers after the cull, b) estimated number of pupal cases and c) colony mass between colonies assigned to a monofloral (intercept) and polyfloral diet on arrival. Asterisks highlight significant differences (alpha values: 0.05 * 0.01 ** 0.001 ***). ER = Experimental Replicate

| a) GLM (worker number ~ diet + ER, family = Poisson) |  |  |  |  |  |
| --- | --- | --- | --- | --- | --- |
|  | Estimate | S.E. | z value | P value |  |
| (Intercept) | 3.875 | 0.054 | 72.139 | <2e-16 | *** |
| dietpoly | 0.023 | 0.068 | 0.339 | 0.735 |  |
| ER1 | -0.718 | 0.072 | -9.955 | <2e-16 | *** |

| b) GLM (pupae number ~ diet + ER, family = Poisson) |  |  |  |  |  |
| --- | --- | --- | --- | --- | --- |
|  | Estimate | S.E. | z value | P value |  |
| (Intercept) | 3.765 | 0.056 | 67.654 | <2e-16 | *** |
| dietpoly | -0.176 | 0.069 | -2.547 | 0.011 | * |
| ER1 | -0.263 | 0.070 | -3.780 | 0.000 | *** |

| c) LM (colony mass ~ diet + ER) |  |  |  |  |  |
| --- | --- | --- | --- | --- | --- |
|  | Estimate | S.E. | t value | P value |  |
| Intercept | 56.864 | 3.467 | 16.402 | 1.9e-13 | *** |
| dietpoly | 5.520 | 4.003 | 1.379 | 0.182 |  |
| ER1 | -16.868 | 4.003 | -4.214 | 0.000 | *** |

**Table S3.**
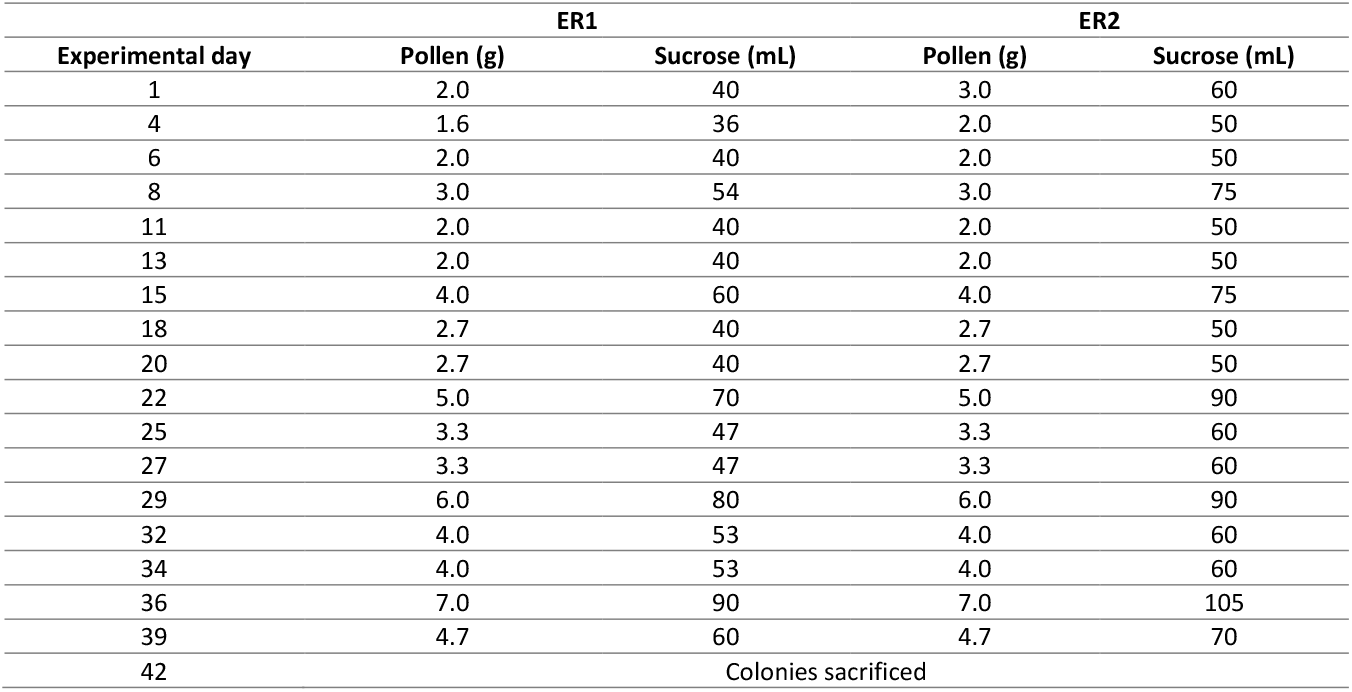
Mass of pollen (g) and volume of 40% sucrose solution provisioned to each colony across both experimental replicates.

| Experimental day | ER1 |  | ER2 |  |
| --- | --- | --- | --- | --- |
|  | Pollen (g) | Sucrose (mL) | Pollen (g) | Sucrose (mL) |
| 1 | 2.0 | 40 | 3.0 | 60 |
| 4 | 1.6 | 36 | 2.0 | 50 |
| 6 | 2.0 | 40 | 2.0 | 50 |
| 8 | 3.0 | 54 | 3.0 | 75 |
| 11 | 2.0 | 40 | 2.0 | 50 |
| 13 | 2.0 | 40 | 2.0 | 50 |
| 15 | 4.0 | 60 | 4.0 | 75 |
| 18 | 2.7 | 40 | 2.7 | 50 |
| 20 | 2.7 | 40 | 2.7 | 50 |
| 22 | 5.0 | 70 | 5.0 | 90 |
| 25 | 3.3 | 47 | 3.3 | 60 |
| 27 | 3.3 | 47 | 3.3 | 60 |
| 29 | 6.0 | 80 | 6.0 | 90 |
| 32 | 4.0 | 53 | 4.0 | 60 |
| 34 | 4.0 | 53 | 4.0 | 60 |
| 36 | 7.0 | 90 | 7.0 | 105 |
| 39 | 4.7 | 60 | 4.7 | 70 |
| 42 | Colonies sacrificed |  |  |  |

**Table S4.**
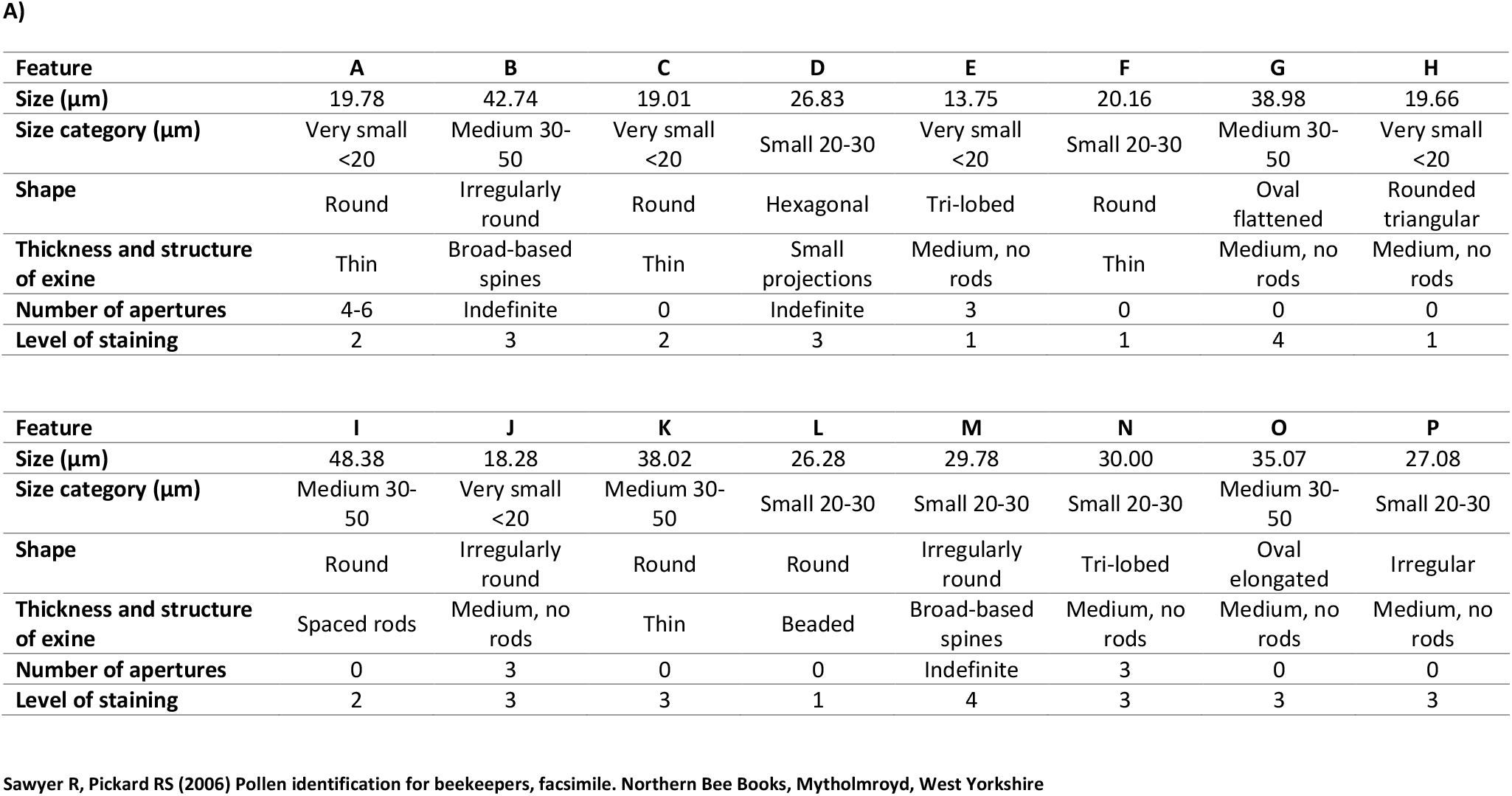
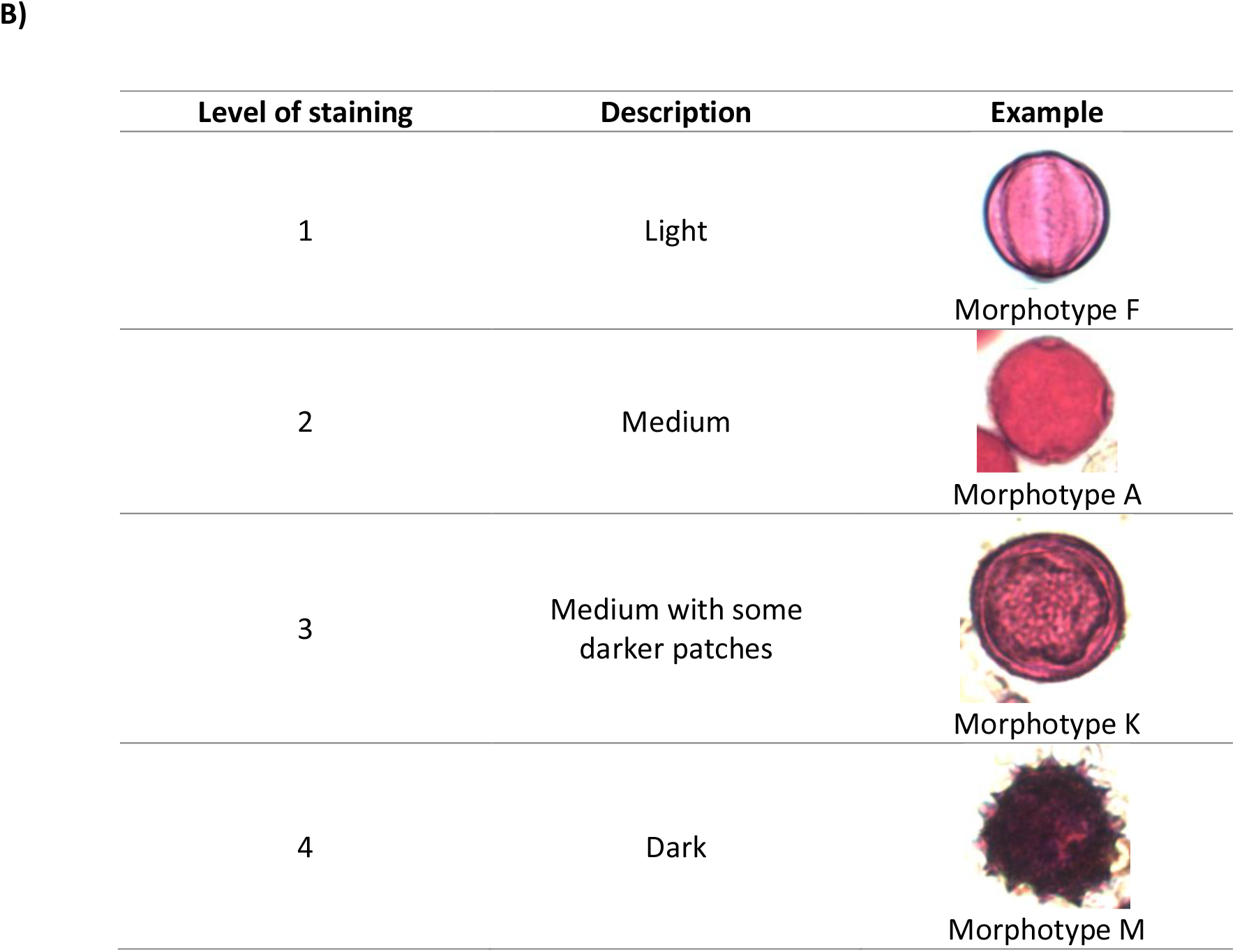
**A)** Descriptions of the 16 pollen morphotypes found across all four diets. Morphotype features are described based on categories outlined in Sawyer (2006): size category (very small <20μm, small 20-30μm, medium 30-50μm, large 50-100μm, very large >100μm), shape, thickness and structure of the exine (outer layer) and number of apertures. Level of staining relates to the level at which the morphotype absorbs the fuchsine dye. **B)** Key used to determine the level of staining of a pollen grain with fuchsine dye, with example morphotypes.

**A)**
| Feature | A | B | C | D | E | F | G | H |
| --- | --- | --- | --- | --- | --- | --- | --- | --- |
| Size (µm) | 19.78 | 42.74 | 19.01 | 26.83 | 13.75 | 20.16 | 38.98 | 19.66 |
| Size category (µm) | Very small<br><20 | Medium 30-50 | Very small<br><20 | Small 20-30 | Very small<br><20 | Small 20-30 | Medium 30-50 | Very small<br><20 |
| Shape | Round | Irregularly<br>round | Round | Hexagonal | Tri-lobed | Round | Oval<br>flattened | Rounded<br>triangular |
| Thickness and structure<br>of exine | Thin | Broad-based<br>spines | Thin | Small<br>projections | Medium, no<br>rods | Thin | Medium, no<br>rods | Medium, no<br>rods |
| Number of apertures | 4-6 | Indefinite | 0 | Indefinite | 3 | 0 | 0 | 0 |
| Level of staining | 2 | 3 | 2 | 3 | 1 | 1 | 4 | 1 |

| Feature | I | J | K | L | M | N | O | P |
| --- | --- | --- | --- | --- | --- | --- | --- | --- |
| Size (µm) | 48.38 | 18.28 | 38.02 | 26.28 | 29.78 | 30.00 | 35.07 | 27.08 |
| Size category (µm) | Medium 30-50 | Very small<br><20 | Medium 30-50 | Small 20-30 | Small 20-30 | Small 20-30 | Medium 30-50 | Small 20-30 |
| Shape | Round | Irregularly<br>round | Round | Round | Irregularly<br>round | Tri-lobed | Oval<br>elongated | Irregular |
| Thickness and structure<br>of exine | Spaced rods | Medium, no<br>rods | Thin | Beaded | Broad-based<br>spines | Medium, no<br>rods | Medium, no<br>rods | Medium, no<br>rods |
| Number of apertures | 0 | 3 | 0 | 0 | Indefinite | 3 | 0 | 0 |
| Level of staining | 2 | 3 | 3 | 1 | 4 | 3 | 3 | 3 |

B)
| Level of staining | Description | Example |
| --- | --- | --- |
| 1 | Light | Morphotype F |
| 2 | Medium | Morphotype A |
| 3 | Medium with some darker patches | Morphotype K |
| 4 | Dark | Morphotype M |

**Table S5.** Composition of each diet by morphotype. Size (μm) was calculated by measuring a single pollen grain of each morphotype relative to a *Lycopodium* spore (33μm at widest point) using the software ImageJ (NIH). The table shows the raw count of pollen grains per morphotype (count), and the percentage compositions wen based on the raw count (%) and when based on the count weighted by grain size (relative %).

| Morph. | Size | Polyfloral pollen diet |  |  |  |  |  | Monofloral pollen diet |  |  |  |  |  |
| --- | --- | --- | --- | --- | --- | --- | --- | --- | --- | --- | --- | --- | --- |
|  |  | ER1 |  |  | ER2 |  |  | ER1 |  |  | ER2 |  |  |
|  |  |  |  | relative |  |  | relative |  |  | relative |  |  | relative |
|  |  | count | % |  | count | % |  | count | % |  | count | % |  |
| A | 19.8 | 37 | 13.6 | 9.1 | 1 | 0.7 | 0.5 | 512 | 97.9 | 96.8 | - | - | - |
| B | 42.7 | 2 | 0.7 | 1.1 | - | - | - | 5 | 1.0 | 2.0 | - | - | - |
| C | 19.0 | - | - | - | - | - | - | 2 | 0.4 | 0.4 | - | - | - |
| D | 26.8 | 8 | 2.9 | 2.7 | 15 | 10.1 | 10.1 | 2 | 0.4 | 0.5 | - | - | - |
| E | 13.8 | - | - | - | 1 | 0.7 | 0.3 | 2 | 0.4 | 0.3 | - | - | - |
| F | 20.2 | 2 | 0.7 | 0.5 | 39 | 26.2 | 19.6 | - | - | - | 412 | 100 | 100 |
| G | 39.0 | 3 | 1.1 | 1.5 | 6 | 4.0 | 5.8 | - | - | - | - | - | - |
| H | 19.7 | 74 | 27.1 | 18.1 | 17 | 11.4 | 8.3 | - | - | - | - | - | - |
| I | 48.4 | 63 | 23.1 | 37.8 | 12 | 8.1 | 14.5 | - | - | - | - | - | - |
| J | 18.3 | 5 | 1.8 | 1.1 | 7 | 4.7 | 3.2 | - | - | - | - | - | - |
| K | 38.0 | 8 | 2.9 | 3.8 | 5 | 3.4 | 4.7 | - | - | - | - | - | - |
| L | 26.3 | 50 | 18.3 | 16.3 | 22 | 14.8 | 14.4 | - | - | - | - | - | - |
| M | 29.8 | 6 | 2.2 | 2.2 | 17 | 11.4 | 12.6 | - | - | - | - | - | - |
| N | 30.0 | 8 | 2.9 | 3.0 | 3 | 2.0 | 2.2 | - | - | - | - | - | - |
| O | 35.1 | 6 | 2.2 | 2.6 | 4 | 2.7 | 3.5 | - | - | - | - | - | - |
| P | 27.1 | 1 | 0.4 | 0.3 | - | - | - | - | - | - | - | - | - |
| Total |  | 273 | 100 | 100 | 149 | 100 | 100 | 523 | 100 | 100 | 412 | 100 | 100 |

**Table S6.** Protein content of the pollen lysate estimated using the Bradford assay for three replicates per diet. SD = standard deviation

|  |  | Protein Content mg/ml |  |  | Mean | SD |
| --- | --- | --- | --- | --- | --- | --- |
|  |  | 1 | 2 | 3 |  |  |
| ER 1 | Polyfloral | 32.290 | 29.102 | 37.408 | 32.933 | 4.190 |
|  | Monofloral | 55.277 | 58.297 | 64.505 | 59.360 | 4.705 |
| ER 2 | Polyfloral | 39.337 | 39.421 | 45.881 | 41.546 | 3.754 |
|  | Monofloral | 58.129 | 58.465 | 72.978 | 63.191 | 8.478 |

**Table S7.** Statistical outputs from linear mixed effects models (LMER) in R for comparing consumption between monofloral (intercept) and polyfloral colonies. **a)** total mass of pollen consumed by colonies per week, **b)** total volume of sucrose solution consumed by colonies per week **c)** mass of pollen consumed per gram of colony mass per week, and **d)**sucrose consumed per gram of colony mass per week. For **a)** and **b)**, a 2^nd^ order polynomial relationship best fitted the data. Asterisks highlighting significant differences (alpha values: 0.05 * 0.01 ** 0.001 ***).

| a) LMER (pollen consumption ~ diet * poly(week,2) + ER + (week\|colony)) |  |  |  |  |  |  |
| --- | --- | --- | --- | --- | --- | --- |
|  | Estimate | S.E. | df | t value | P value |  |
| (Intercept) | 9.583 | 0.454 | 22.462 | 21.111 | 2.62e-16 | *** |
| dietpoly | -0.160 | 0.548 | 19.714 | -0.293 | 0.773 |  |
| poly(week, 2) 1 | 21.325 | 2.735 | 21.993 | 7.798 | 9.02-e08 | *** |
| poly(week, 2) 2 | -11.159 | 2.006 | 94.000 | -5.562 | 2.49e-07 | *** |
| ER1 | -1.765 | 0.474 | 21.000 | -3.725 | 0.001 | ** |
| dietpoly:poly(week,2) 1 | -1.735 | 3.867 | 21.993 | -0.449 | 0.658 |  |
| dietpoly:poly(week,2) 2 | 0.125 | 2.837 | 94.000 | 0.044 | 0.965 |  |

| b) LMER (sucrose consumption ~ diet * poly(week,2) + ER + (week\|colony)) |  |  |  |  |  |  |
| --- | --- | --- | --- | --- | --- | --- |
|  | Estimate | S.E. | df | t value | P value |  |
| (Intercept) | 169.950 | 4.289 | 21.587 | 39.628 | <2e-16 | *** |
| dietpoly | -20.239 | 5.165 | 18.552 | -3.918 | 0.001 | *** |
| poly(week,2)1 | 128.272 | 35.922 | 21.999 | 3.571 | 0.002 | ** |
| poly(week,2)2 | -109.746 | 24.165 | 93.998 | -4.541 | 1.66e-05 | *** |
| ER1 | -46.161 | 4.496 | 21.000 | -10.267 | 1.22e-09 | *** |
| dietpoly:poly(week,2)1 | -95.765 | 50.801 | 21.999 | -1.885 | 0.073 |  |
| dietpoly:poly(week,2)2 | 32.904 | 34.175 | 93.998 | 0.963 | 0.338 |  |

| c) LMER (pollen consumed per gram colony mass ~ diet * week + (week\|colony)) |  |  |  |  |  |  |
| --- | --- | --- | --- | --- | --- | --- |
|  | Estimate | S.E. | df | t value | P value |  |
| (Intercept) | 1.036 | 0.307 | 22.012 | 3.376 | 0.003 | ** |
| dietpoly | 0.079 | 0.434 | 22.012 | 0.181 | 0.858 |  |
| week | -0.105 | 0.037 | 22.010 | -2.794 | 0.011 | * |
| dietpoly:week | -0.007 | 0.053 | 22.010 | -0.125 | 0.902 |  |

| d) LMER (sucrose consumed per gram colony mass ~ diet * week + (week\|colony)) |  |  |  |  |  |  |
| --- | --- | --- | --- | --- | --- | --- |
|  | Estimate | S.E. | df | t value | P value |  |
| (Intercept) | 2.733 | 0.103 | 22.000 | 26.636 | <2e-16 | *** |
| dietpoly | -0.109 | 0.145 | 22.000 | -0.752 | 0.460 |  |
| week | -0.198 | 0.024 | 22.000 | -8.210 | 3.83e-08 | *** |
| dietpoly:week | 0.017 | 0.034 | 22.000 | 0.510 | 0.615 |  |

**Table S8.** Statistical output from a linear mixed effects model (LMER) in R comparing cumulative increase in colony mass between monofloral (intercept) and polyfloral colonies. A 2^nd^ order polynomial relationship best fitted the data, and asterisks highlight significant differences (alpha values: 0.05 * 0.01 ** 0.001 ***).

| LMER (cumulative colony mass ~ diet * poly(week,2) + ER + (week\|colony)) |  |  |  |  |  |  |
| --- | --- | --- | --- | --- | --- | --- |
|  | Estimate | S.E. | df | t value | P value |  |
| (Intercept) | 81.140 | 3.435 | 20.848 | 23.625 | <2e-16 | *** |
| dietpoly | -6.354 | 4.189 | 17.111 | -1.517 | 0.148 |  |
| poly(week,2)1 | 250.121 | 19.427 | 22.000 | 12.875 | 1.02e-11 | *** |
| poly(week,2)2 | 23.402 | 6.494 | 118.001 | 3.604 | 0.000 | *** |
| ER1 | -18.320 | 3.478 | 20.998 | -5.268 | 3.19e-05 | *** |
| dietpoly:poly(week,2)1 | -88.658 | 27.474 | 22.000 | -3.227 | 0.004 | ** |
| dietpoly:poly(week,2)2 | 13.262 | 9.183 | 118.001 | 1.444 | 0.151 |  |

**Table S9.** Statistical output from a generalised linear mixed effects model (GLMER) in R comparing cumulative increase in worker production (reared) between monofloral (intercept) and polyfloral colonies **a)** excluding starting pupae number and **b)** including starting pupae number as a fixed factor in the model. A 2^nd^ order polynomial relationship best fitted the data, and asterisks highlight significant differences (alpha values: 0.05 * 0.01 ** 0.001 ***). For model b), we included starting pupae number to account for polyfloral colonies arriving with a significantly lower number of pupae. However, running the model presented convergence warnings. Furthermore, it did not significantly alter the results of the model, and so we presented only model a) in the main text.

| a) GLMER (colony_worker_growth ~ diet * poly(week,2) + ER + (week\|colony), family = Poisson) |  |  |  |  |  |
| --- | --- | --- | --- | --- | --- |
|  | Estimate | S.E. | z value | P value |  |
| (Intercept) | 4.740 | 0.107 | 44.395 | <2e-16 | *** |
| dietpoly | -0.199 | 0.122 | -1.635 | 0.102 |  |
| poly(week,2)1 | 6.614 | 0.475 | 13.926 | <2e-16 | *** |
| poly(week,2)2 | -1.243 | 0.149 | -8.328 | <2e-16 | *** |
| ER1 | -0.429 | 0.127 | -3.389 | 0.001 | *** |
| dietpoly:poly(week,2)1 | -0.122 | 0.678 | -0.180 | 0.857 |  |
| dietpoly:poly(week,2)2 | -0.052 | 0.219 | -0.236 | 0.813 |  |

| b) GLMER (colony_worker_growth ~ diet * poly(week,2) + ER + starting pupae number + (week\|colony), family = Poisson) |  |  |  |  |  |
| --- | --- | --- | --- | --- | --- |
|  | Estimate | S.E. | z value | P value |  |
| (Intercept) | 4.072 | 0.184 | 22.121 | <2e-16 | *** |
| dietpoly | -0.098 | 0.097 | -1.013 | 0.311 |  |
| poly(week,2)1 | 6.615 | 0.472 | 14.009 | <2e-16 | *** |
| poly(week,2)2 | -1.244 | 0.149 | -8.338 | <2e-16 | *** |
| ER1 | -0.311 | 0.101 | -3.080 | 0.002 | ** |
| starting pupae number | 0.016 | 0.004 | 4.057 | 4.97e-05 | *** |
| dietpoly:poly(week,2)1 | -0.157 | 0.674 | -0.233 | 0.816 |  |
| dietpoly:poly(week,2)2 | -0.047 | 0.219 | -0.216 | 0.829 |  |

**Table S10.** Statistical outputs from generalised linear models (GLM) in R when analysing the difference in total number of a) and b) workers and c) pupae by the end of the experiment for colonies assigned to monofloral (intercept) and polyfloral diet. Asterisks highlight significant differences (alpha values: 0.05 * 0.01 ** 0.001 ***). For model b), we again included starting pupae number in the model analysing total worker production, in order to account for polyfloral colonies arriving with a significantly lower number of pupae. Including starting pupae number did not appear to affect the result of the models, and so we presented only model a) in the main text. Note that model c) excludes data for Colony D, which had begun producing gyne pupae (~600 mg) and was therefore not included in the analysis.

|  | Estimate | S.E. | z value | P value |  |
| --- | --- | --- | --- | --- | --- |
| (Intercept) | 5.372 | 0.025 | 215.586 | <2e-16 | *** |
| dietpoly | -0.192 | 0.031 | -6.170 | 6.8e-10 | *** |
| ER1 | -0.253 | 0.031 | -8.135 | 4.11e-16 | *** |

b) GLM (workers ~ diet + ER + starting pupae number, family = Poisson)
|  | Estimate | S.E. | z value | P value |  |
| --- | --- | --- | --- | --- | --- |
| (Intercept) | 4.921 | 0.058 | 85.451 | <2e-16 | *** |
| dietpoly | -0.134 | 0.032 | -4.224 | 2.39e-05 | *** |
| ER1 | -0.150 | 0.034 | -4.457 | 8.30e-06 | *** |
| starting pupae number | 0.010 | 0.001 | 8.924 | <2e-16 | *** |

c) GLM (pupae ~ diet + ER, family = Poisson)
|  | Estimate | S.E. | z value | P value |  |
| --- | --- | --- | --- | --- | --- |
| (Intercept) | 4.437 | 0.042 | 106.994 | <2e-16 | *** |
| dietpoly | -0.189 | 0.056 | -3.385 | 0.001 | *** |
| ER1 | -0.832 | 0.061 | -13.556 | <2e-16 | *** |

**Table S11.** Statistical output from a linear mixed effects model (LMER) in R comparing change in worker mass over the course of the experiment between monofloral (intercept) and polyfloral colonies. A 2^nd^ order polynomial relationship best fitted the data, and asterisks highlight significant differences (alpha values: 0.05 * 0.01 ** 0.001 ***).

| LMER (worker mass ~ diet * poly(week,2) + ER + (week\|colony)) |  |  |  |  |  |  |
| --- | --- | --- | --- | --- | --- | --- |
|  | Estimate | S.E. | df | t value | P value |  |
| (Intercept) | 130.653 | 5.526 | 19.849 | 23.644 | 5.21e-16 | *** |
| dietpoly | -10.105 | 6.351 | 19.620 | -1.591 | 0.128 |  |
| poly(week,2)1 | -814.121 | 190.881 | 21.758 | -4.265 | 0.000 | *** |
| poly(week,2)2 | 789.219 | 48.318 | 3334.620 | 16.334 | <2e-16 | *** |
| ER1 | 15.340 | 6.356 | 19.636 | 2.413 | 0.026 | * |
| dietpoly:poly(week,2)1 | -728.213 | 270.166 | 22.087 | -2.695 | 0.013 | * |
| dietpoly:poly(week,2)2 | 37.307 | 73.800 | 3341.534 | 0.506 | 0.613 |  |

**Table S12.**
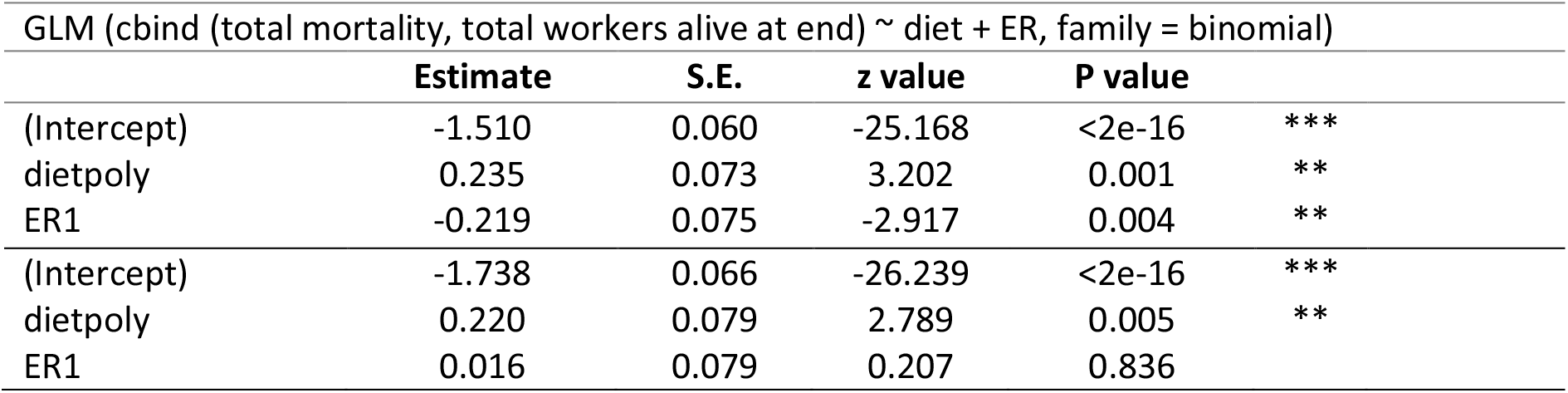
Statistical outputs from generalised linear models (GLM) in R when analysing the difference in total mortality of workers by the end of the experiment between monofloral (intercept) and polyfloral diet for colonies that included (top three rows) and excluded (lower three rows) the two colonies (one monofloral, one polyfloral) experiencing >50% mortality. Asterisks highlight significant differences (alpha values: 0.05 * 0.01 ** 0.001 ***).

**Table S13.** Total number of males produced by each colony across experimental replicates 1 and 2.

| Experimental replicate | Colony identification | Diet | Total number of males produced |
| --- | --- | --- | --- |
| 1 | A | Polyfloral | 0 |
| 1 | B | Polyfloral | 33 |
| 1 | C | Polyfloral | 0 |
| 1 | D | Polyfloral | 0 |
| 1 | E | Polyfloral | 0 |
| 1 | F | Polyfloral | 0 |
| 1 | G | Monofloral | 6 |
| 1 | H | Monofloral | 3 |
| 1 | I | Monofloral | 1 |
| 1 | J | Monofloral | 0 |
| 1 | K | Monofloral | 1 |
| 1 | L | Monofloral | 0 |
| 2 | AA | Monofloral | 3 |
| 2 | BB | Monofloral | 45 |
| 2 | CC | Monofloral | 7 |
| 2 | DD | Monofloral | 108 |
| 2 | EE | Monofloral | 32 |
| 2 | FF | Monofloral | 5 |
| 2 | GG | Polyfloral | 7 |
| 2 | HH | Polyfloral | 4 |
| 2 | II | Polyfloral | 1 |
| 2 | JJ | Polyfloral | 1 |
| 2 | KK | Polyfloral | 47 |
| 2 | LL | Polyfloral | 15 |

**Table S14.** Mean (±s.d.) number of larvae and pupae found per colony on dissection at the end of the experiment, as well as mass (g) of the nest structure once all brood, adult workers and the queen had been removed.

|  | Monofloral diet | Polyfloral diet |
| --- | --- | --- |
| Small larvae | 26.3 ± 7.0 | 19.4 ± 4.3 |
| Medium larvae | 28.8 ± 9.0 | 29.3 ± 4.9 |
| Large larvae | 5.3 ± 1.3 | 5.3 ± 1.5 |
| Pupae | 60.7 ± 11.2 | 49.3 ± 9.3 |
| Nest structure (g) | 20.6 ± 1.6 | 21.1 ± 2.4 |

**Table S15.** Statistical outputs from linear mixed effects models (LMER) in R when analysing **a)** male mass and **b)** average pupal mass per colony dissected from the nest, at the end of the experiment between monofloral (intercept) and polyfloral colonies. For male mass the data was log10 transformed to better meet the assumptions of normality. For the pupal mass, one polyfloral colony was not considered in this analysis because it was producing gyne pupae (~600mg) at the end of the experiment. Asterisks highlight significant differences (alpha values: 0.05 * 0.01 ** 0.001 ***).

| a) LMER (male mass ~ diet + ER + (1\|colony)) |  |  |  |  |  |
| --- | --- | --- | --- | --- | --- |
|  | Estimate | S.E. | df | t value | P value |
| (Intercept) | 2.326 | 0.039 | 8.727 | 59.673 | 1.05e-12 *** |
| dietpoly | -0.113 | 0.054 | 9.607 | -2.103 | 0.063 |
| ER1 | -0.022 | 0.064 | 10.975 | -0.344 | 0.737 |

| b) LM (pupa mean mass ~ diet + ER) |  |  |  |  |
| --- | --- | --- | --- | --- |
|  | Estimate | S.E. | t value | P value |
| (Intercept) | 334.24 | 25.65 | 13.032 | 3.12e-11 *** |
| dietpoly | -73.48 | 30.08 | -2.410 | 0.026 * |
| ER1 | 19.73 | 30.08 | 0.656 | 0.519 |

